# Different CMV-specific effector T cell subtypes are associated with age, CMV serostatus, and increased systolic blood pressure

**DOI:** 10.1101/2024.07.26.605265

**Authors:** LM Roesner, S Traidl, B Bošnjak, J Huehn, R Förster, T Werfel

## Abstract

Cytomegalovirus (CMV) infection is one of the most common infections in humans, and CMV antigens are the major drivers of repetitive T-cell stimulation as a part of a well-adapted immune response in immunocompetent individuals. With higher age, the recurrent clonal expansion of CMV-specific T cells results in high frequencies of CMV-specific effector T cells. CMV seropositivity has also been linked to arterial stiffness and an increased risk of developing cardiovascular diseases (CVD). The RESIST Senior Individuals (SI) cohort is a population-based cohort with focus on the elderly, established to shed light on the age-related changes of the immune system and the accompanied reduced capability to fight infectious diseases.

Here we investigated the frequency and phenotype of CMVpp65-specific CD8^+^ T cells in the circulation of individuals of different age groups by means of MHC-I tetramer staining and their associations with age and associated factors such as serostatus and blood pressure.

In the SI cohort, the frequency of CMV-specific T cells within the CD8^+^ T cell fraction was increased with age, as previously reported. We add to previous knowledge by showing that this frequency is associated with the total percentage and absolute counts of CD8^+^ and CD4^+^CD8^+^ double-positive T cells within leukocytes. Systolic blood pressure (SBP) and history of CVD correlated with the frequency of CMV-specific CD8^+^ T cells. Focusing on CMV-specific T cell subtypes, we show here that the frequencies of T_EM_ and CD27-expressing T_EMRA_ cells were associated with higher age. T_EM_ and CD27^-^ T_EMRA_ cell frequencies were increased in donors with high CMV-IgG titers. Furthermore, SBP significantly correlated with CMV-specific effector CD8^+^ T cells, which was mostly reflected by CD27^-^ T_EMRA_ cells.

In conclusion, different effector T-cell subtypes were associated with age, serostatus and SBP, suggesting that it is not age *per se* that renders elderly CMV-positive individuals susceptible to CVD, but the immune response to CMV. Our study suggests that detailed immunophenotyping may identify individuals whose immune systems are strongly influenced by the response to CMV, leading to health consequences and impairing healthy aging.

## Introduction

Aging is a progressive biological process in most multicellular organisms that leads to the gradual loss of healthy body and organ functions and ultimately to biological death. The immune system undergoes significant changes during aging, resulting in a diminished acquired immunity accompanied by an increased inflammatory phenotype [1]. As a result of thymic involution, there are measurably fewer naive T cells in the blood and secondary lymphoid organs compared to younger adults [2]. But thymic involution does not represent a crucial loss for the immune system. Even if the activity level is not comparable to the fetal and perinatal period, new naive T cells are exported into old age [3]. The total number of T cells is nearly maintained by a number of thymus-independent homeostatic mechanisms: Long-lived naive T cells proliferate at a low rate in response to low affinity antigens, and memory T cells undergo continuous cytokine-driven proliferation [4]. The overweight of memory T cells is also a consequence of the interaction of naive cells with environmental antigens. Newly activated naive and memory cell populations expand in response to cognate antigen stimulation and occupy larger portions of the repertoire (e.g., after infection), resulting in a relative increase of memory cell frequencies in older people, but also to a smaller repertoire, but no general defects in the adaptive immune system. In particular, infection with herpesviruses that persist lifelong in the organism undergoing latency with continuous reactivation, such as Cytomegalovirus (CMV), lead to a large number of specific memory cells [5]. In CMV-positive elderly individuals, CMV-specific CD8^+^ T cells can account for up to a quarter of the total CD8^+^ T cell population [6], which may lead to a reduced ability to respond to novel threats in the elderly [7, 8], but not in young adults [9]. This accumulation of CMV-specific CD8+ T cells exhausts the immune system over time and accelerates immunosenescence [10]. Therefore, latent CMV infection has the potential to influence the course of further infections, due to the broader reshaping of cytotoxic lymphocyte populations. On the other hand, CMV infection has been linked to an increased risk towards cardiovascular diseases [11-13] and thereby also towards death in elderlies [14], reflected also by an increased systolic blood pressure [15].

Because of the strong effects, many population or cohort studies examining the impact of CMV on the immune response are based on a comparison between seropositive and negative individuals [16]. To investigate these effects in with a focus on the cellular response, we investigated CMV-specific T cell frequencies and phenotypes in the RESIST senior individuals (SI) cohort. This population-based cohort of 650 citizens with a focus on the age group of 60-100 years was set up between 2019 and 2023 to investigate age-related changes of the immune system. A combination of clinical data, immune phenotyping and multi-omics analyzes build the core of the study [17]. Here, we report on frequency of CMV-specific T cells from fresh blood and their differentiation status with regard to age, sex, antibody titer, blood pressure, and patient-reported history of cardiovascular diseases.

## Methods

### Patients and Ethics

The SI cohort is was conducted in compliance with all pertinent legislation and directives and following the guidelines on human biobanks for research and other relevant directives on research ethics. Special emphasis is placed on privacy, safety of data, genetic information and reporting of incidental medical findings to participants. Ethics committee of Hannover Medical School (MHH), Hannover, Germany, gave ethical approval for this work (No 8615_BO_S_2019; connected disease cohorts under the file references 8730_BO_S_2019 and 8733_BO_S_2019). The study was conducted according to the principles of declaration of Helsinki. Participants gave their written informed consent prior to the study.

### Antibody titer testing

Blood was drawn without anticoagulant for serum sampling and with EDTA for PBMC isolation. Serum samples were tested in the routine diagnostic of the clinical virology lab at MHH. Serology was performed on the Architect System from Abbott Diagnostics (Abbott GmbH & Co. KG, Wiesbaden, Germany) using the CMV IgG assay for detection of IgG antibodies against CMV. The Detection of IgG antibodies for VZV and HSV was done on the Liaison XL analyzer from Diasorin (DiaSorin S.p.A. Saluggia (VC), Italy) using the LIAISON® VZV IgG and the LIAISON® HSV-1/2 IgG assay.

### Immune cell count

Whole fresh blood was subjected to FACS Lysing Solution (#349202, BD Biosciences, Franklin Lakes, NJ, USA) and subsequently the 6-color TBNK Reagent with TruCount tubes (#337166, BD Biosciences) according to the manufacturer’s protocol (Supplemental Figure 2). Fluorescence signals and TruCount beads were acquired on a 3-laser CytoFLEX using Cytexpert software 2.3 (Beckman Coulter, Brea, CA, USA). Instrument quality control and standardization were performed daily using CytoFLEX Daily QC Fluorospheres (Beckman Coulter #B53230).

### Analysis of CMV-specific T cells

HLA-A2 status was determined by antibody staining (clone BB7.2, BD Biosciences) on CD45^+^ cells (clone HI30, BioLegend, San Diego, CA, USA) in fresh blood after subjecting to FACS Lysing Solution (BD Biosciences) by means of flowcytometry. PBMC were isolated by density-gradient centrifugation on Ficoll (Pan Biotech, Aidenbach, Germany) and subjected to subsequent analyses. 1×10^6^ cells were incubated with MHC-I-tetramers, consisting of tetramerized and PE-labelled HLA-A*02:01 monomers carrying the NLVPMVATV peptide CMVpp65_495-503_ (1002-07 ImmunAware, Horsholm, Denmark; IEDB#: 44920) at room temperature for 30min. Subsequently, anti-CD8-BV510 (clone SK1, Biolegend 344732), anti-CD27-APC (clone O323, Biolegend 302810), anti-CD45RO-PacificBlue (clone UCHL1, Biolegend 304216) and a viability dye (ThermoFisher/Invitrogen/eBioscience fixable viability dye eFluor780) were added and incubated for 30min at 4°C. After washing, cells were analyzed on aforementioned 3-laser CytoFLEX using Cytexpert software 2.3. Data were analyzed with the Software Kaluza V1.3, BeckmanCoulter.

### Statistical analyses

We used GraphPad Prism (Version 5.0.0, GraphPad, La Jolla, USA) and R (version 4.1.2) with the gt-summary package (version 1.7.0) for statistical analyses using tests as listed under each figure. The Wilcoxon matched pairs test was applied when comparing two groups. The Kruskal-Wallis test with Dunn’s multiple comparison test was used for comparing more than two independent samples. Spearman’s rank correlation was applied to measure the strength and direction of association between two ranked variables. Generalized linear mixed models (GLMM) were applied to adjust for age, sex and body mass index. Results with a p-value < 0.05 were considered statistically significant (*p < 0.05; **p < 0.01; ***p < 0.001).

## Results

### Stratification of participants of the RESIST SI cohort

In total, 650 individuals with focus on the elderly were recruited in the RESIST SI cohort and followed a structured study visit as depicted elsewhere [17]. During the first study visit, we analyzed HLA-A*02 expression on blood cells using flow cytometry if there was enough material available for the follow-up examinations on CMV-specific T cells. Of the 596 analyzed samples, 51% (n=302) stained positive for HLA-A*02 (Figure 1A). Next, MHC-I tetramers carrying the CMVpp65 peptide epitope NLVPMVATV were used to analyze the frequency of CMV-specific T cells within the T cell fraction, and antibodies against CD45RO and CD27 were used in parallel to investigate their differentiation status (Figure 1B).

**Figure 1.**
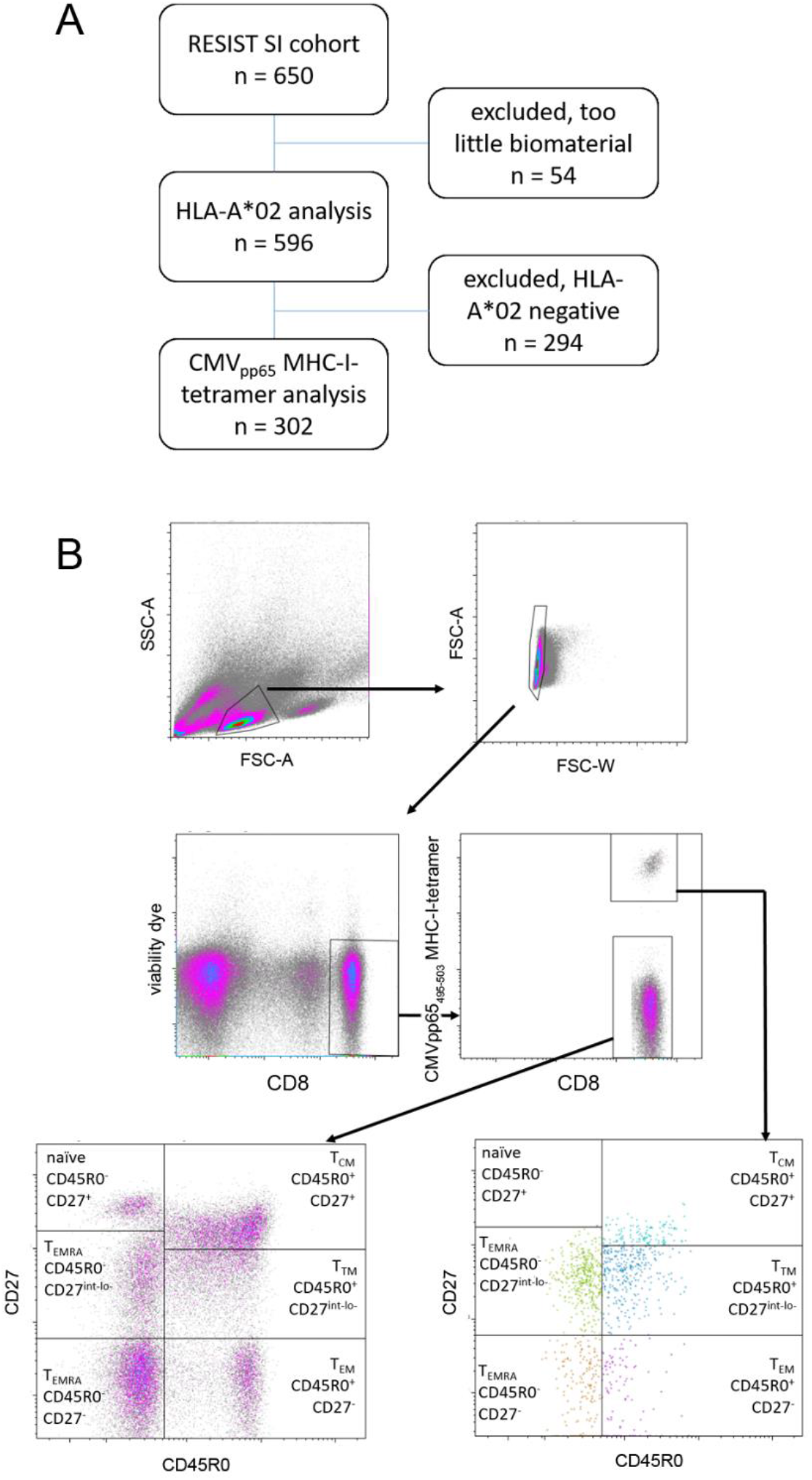
Study outline. A, From the 650 healthy volunteers of the RESIST senior individuals (SI) cohort, a total of 302 individuals was analyzed by means of CMVpp65_495-503_ MHC-I-tetramer staining. B, flow cytometry gating scheme discriminating the differentiation status of the CMV-specific T cells. Vital, singlet CD8^+^ lymphocytes of HLA-A*02 positive individuals were incubated with PE-labelled CMVpp65_495-503_ MHC-I-tetramers and antibodies to CD27 and CD45RO to subtype CMV-specific T cells into naive T (CD45RO^-^CD27^hi^), T_CM_ (CD45RO^+^CD27^+^), T_TM_ (CD45RO^+^CD27^int^), and three effector subtypes: T_EM_ (CD45RO^+^CD27^-^), CD45RO^-^CD27^-^ T_EMRA_ and CD45RO^-^CD27^int-lo^ T_EMRA._

### Higher frequencies of CMV-specific CD45RO^+^/CD27^-^ and CD45RO^-^/CD27^lo-int^ in elderlies

Frequencies of CMV-specific T cells comparing young adults (20-40 years-old) and elderly (60 years or older) individuals are given in Figure 2A,B. Confirming earlier reports, the frequency was increased in the elderlies, while no significant differences were to be seen when comparing male and female participants (Supplemental Figure 1). It is well-established, that naive T cells highly express CD27 while lacking expression of CD45RO. CD27 expression is gradually lost when T cells differentiate into T_CM_, T_TM_, T_EM_, and terminally differentiated T_EMRA_ (or: T_TE_ cells) [18]. Interestingly, the latter do mostly but not always lack CD27 [19-21]. In our approach, we identified the T naive (CD45RO^-^CD27^hi^), T_CM_ (CD45RO^+^CD27^+^), T_TM_ (CD45RO^+^CD27^int^), T_EM_ (CD45RO^+^CD27^-^), as well as CD45RO^-^CD27^-^ and CD45RO^-^ CD27^int-lo^ T_EMRA_ cells within the CMVpp65 MHC-I-tetramer^+^ cells and investigated the frequencies in relation to the factor age (Figure 2C and D). The CMVpp65-specific T_TM_ as well as the rare specific naive T cells were found to be decreased in the elderlies compared to the young adults. On the other hand, the T effector fraction of CMVpp65-tetramer^+^ cells, comprising the T_EM_ and the T_EMRA_ subtypes, increased, while T_CM_ frequencies were not significantly different. Applying a finer categorization, the T cell subtypes found to be increased carried the marker combinations CD45RO^-^CD27^-^ and CD45RO^-^ CD27^int-lo^. These differences were also visible by trend when comparing the elderly participants in age-groups of five years (Supplemental Figure 2).

**Figure 2.**
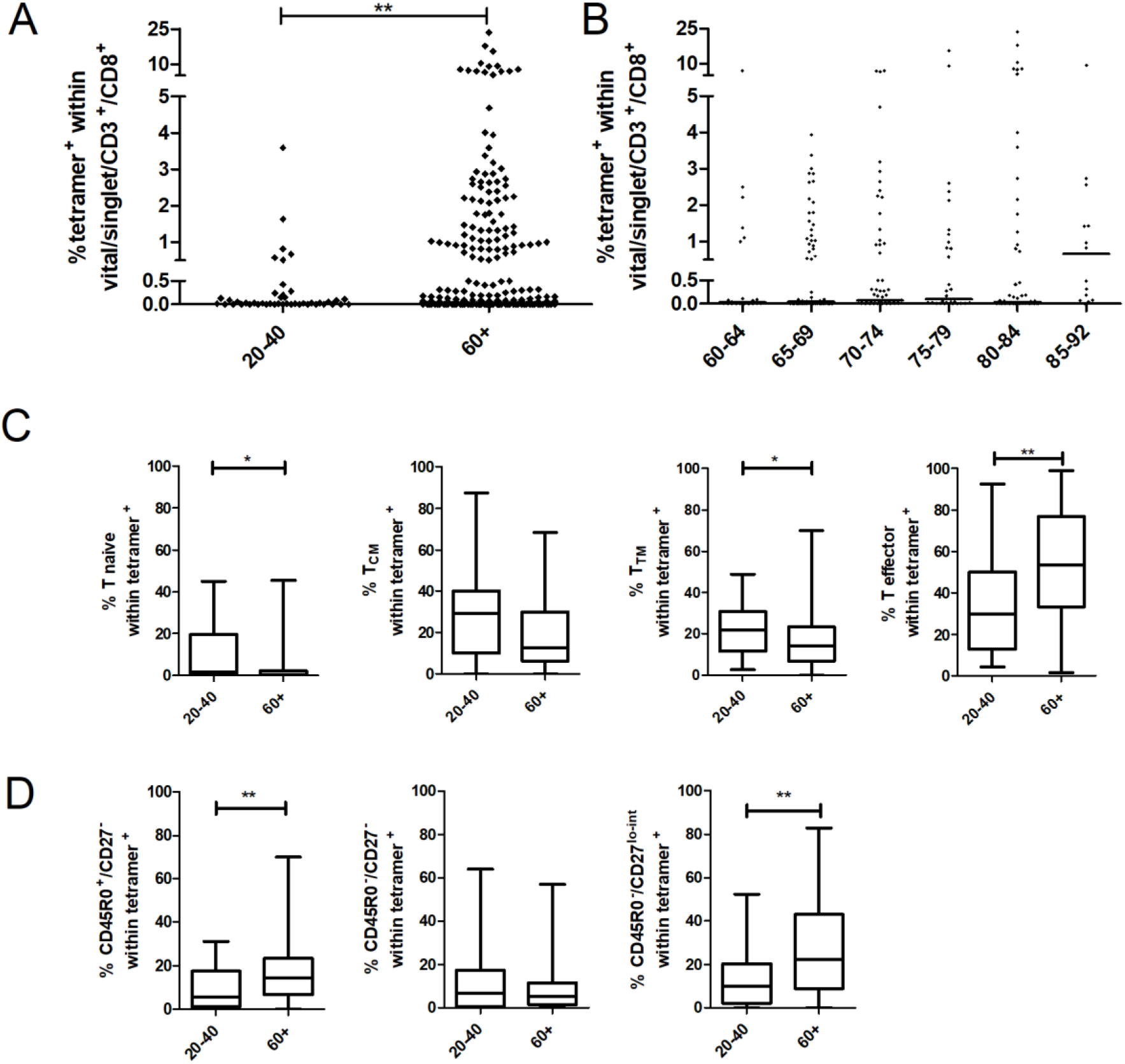
Frequency and differentiation status of CMV-specific T cells with regard to age. A, Frequency of CMVpp65_495-503_ MHC-I-tetramer^+^ T cells comparing the age groups of 20-40 and 60+ year old healthy volunteers. B, Frequency of CMVpp65_495-503_ MHC-I-tetramer^+^ T cells among the 60+ year old healthy volunteers, dissected into groups of 5 years of age. C, Frequency of naïve, TCM, TTM, and effector T cells among CMVpp65_495-503_ MHC-I-tetramer^+^ T cells comparing the age groups of 20-40 and 60+ year old healthy volunteers. D, CMVpp65_495-503_ MHC-I-tetramer^+^ T effector T cells were further dissected into T_EM_, CD27^-^ T_EMRA_ and CD27^int^ T_EMRA_ groups as indicated. Wilcoxon matched pairs test, *p < 0.05; **p < 0.01.

### CMV-specific IgG is associated with higher frequencies of CMV-specific T_EM_ and CD45RO^-^/CD27^-^ T_EMRA_

The individual humoral response to CMV was assessed by clinical routine serum analytics. No striking differences were observed comparing young adults and elderlies regarding the CMV-IgG titers, but elderly donors without detectable CMVpp65-specific T cells differed from those with a cellular response (Supplemental Figure 3). Higher frequencies of CMVpp65-specific T cells were detectable in donors with high antibody titers (Figure 3A). Dissecting these T cells regarding their differentation status, a reduction in naive and T_CM_ and an increase of the T effector fraction were observed comparing individuals with high and low CMV-IgG titers (Figure 3B). More precise, both the T_EM_ and CD45RO^-^ /CD27^-^ T_EMRA_ subtype frequencies were increased (Figure 3C). No association was observed between the CMV-specific T cell frequency and Herpes simplex virus (HSV)- and Varicella zoster virus (VZV)-IgG titers (Supplemental Figure 4).

**Figure 3.**
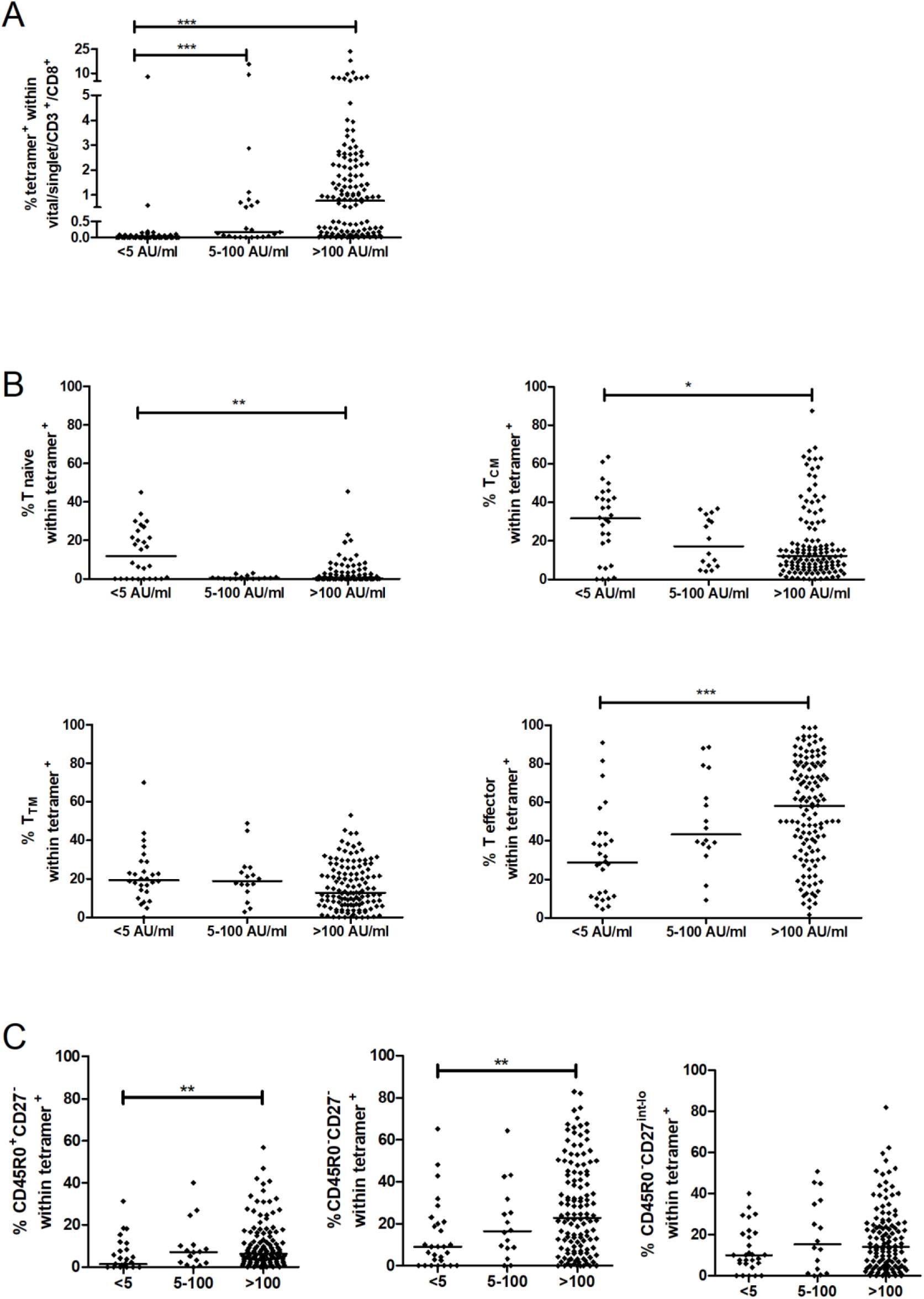
Frequency and differentiation status of CMV-specific T cells with regard to anti-CMV-IgG titers. Healthy volunteers were grouped according to the serum titers of anti-CMV-IgG (AU/ml) as indicated. A, Frequency of CMVpp65_495-503_ MHC-I-tetramer^+^ T cells. B, Frequency of naïve, T_CM_, T_TM_, and effector T cells among CMVpp65_495-503_ MHC-I-tetramer^+^ T cells. C, CMVpp65_495-503_ MHC-I-tetramer^+^ T effector T cells were further dissected into T_EM_, CD27^-^ T_EMRA_ and CD27^int^ T_EMRA_ groups as indicated. Kruskal-Wallis test with Dunn’s multiple comparison testing, *p < 0.05; **p < 0.01; ***p < 0.001.

### High frequencies of CMV-specific T cells correlate with overall CD8^+^ and CD4+CD8+ T cell frequencies and absolute cell counts

In line with previous reports, CMVpp65-specific T cells varied strongly between individuals, with frequencies often exceeding 0.5% of total CD8^+^ T cells. To investigate whether these percentages were also reflected in the composition of the blood count, we compared the data with the absolute leukocyte counts. Individuals were assigned regarding the frequency of CMVpp65 MHC-I-tetramer^+^ cells within the CD8^+^ fraction to a low (<0.5%), a medium (0.5-2%), and a high (>2%) group. Individuals of the low group had significantly less total CD8^+^ T cell and CD4^+^CD8^+^ double-positive T cell frequencies compared to the medium and the high group (Figure 4A). Of note, this was mostly reflected by higher absolute counts of CD8^+^ and CD4^+^CD8^+^ double-positive T cells (Figure 4B). Since also for the factors age, sex, and body mass index an impact on the composition of the blood count has been described, we applied generalized mixed models to exclude confounding effects. Figure 4C shows that CMV-specific T-cell frequency is responsible for the effects, while age and sex partially confound the different parameters and blur the effects.

**Figure 4.**
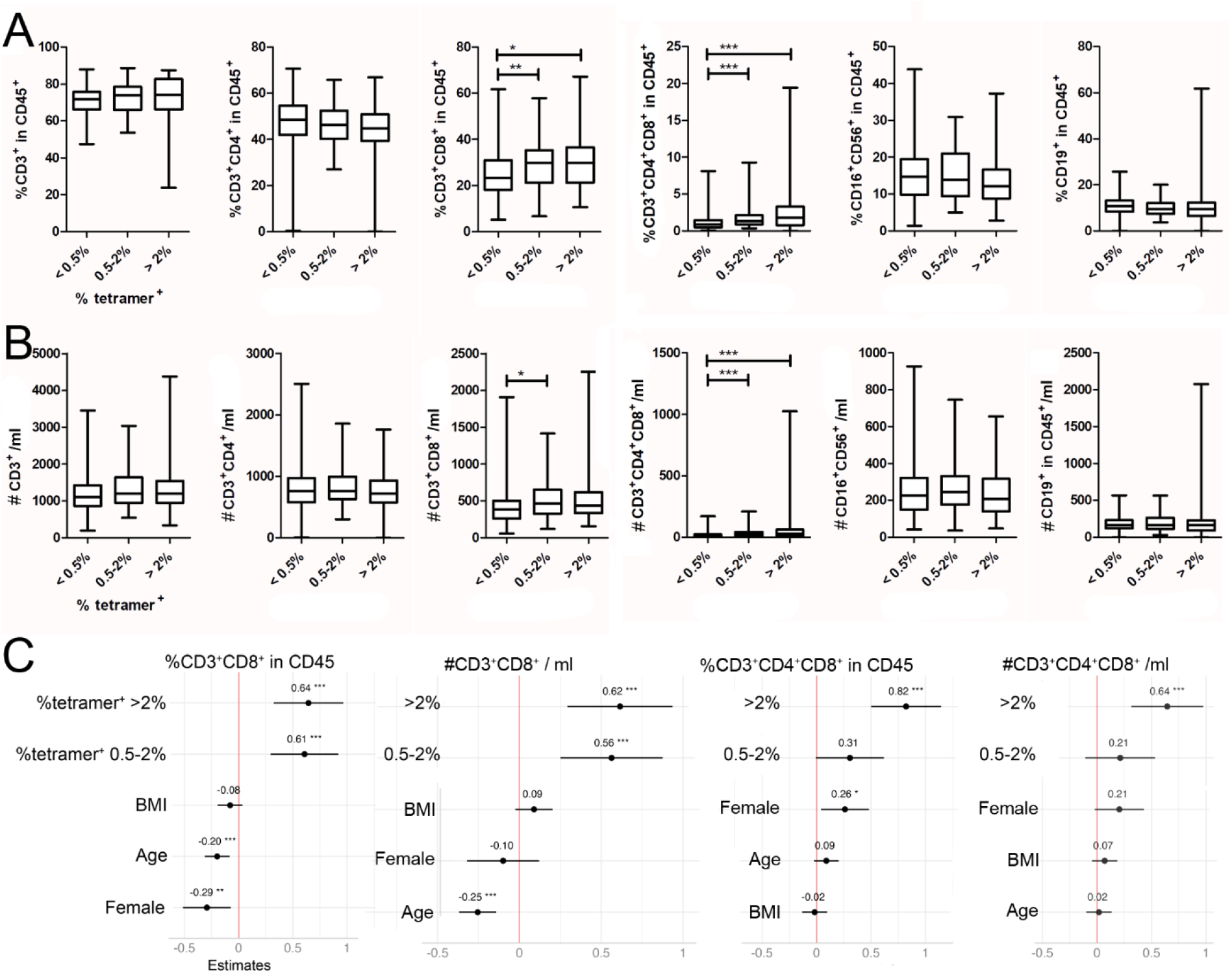
Association of the frequency of CMV-specific T cells and total leukocyte frequencies and counts. Subjects were grouped according to the frequency of CMVpp65_495-503_ MHC-I-tetramer^+^ T cells within the vital CD3+CD8+ fraction as indicated into <0.5%, 0.5-2%, and >2%. A, Frequencies of T cells (CD3^+^), CD3^+^CD4^+^, CD3^+^CD8^+^, CD3^+^CD4^+^CD8^+^ double positive T cells, NK cells (CD16^+^CD56^+^), and B cells (CD19^+^) as determined by TruCount flow cytometry in fresh blood. B, Absolute counts of T cells (CD3^+^), CD3^+^CD4^+^, and CD3^+^CD8^+^, CD3^+^CD4^+^CD8^+^ double positive T cells, NK cells (CD16^+^CD56^+^), and B cells (CD19^+^) as determined by TruCount flow cytometry in fresh blood. Kruskal-Wallis test with Dunn’s multiple comparison testing. C, Forest plots of generalized mixed models. Total frequencies and counts of CD3^+^CD8^+^ and CD3^+^CD4^+^CD8^+^ double positive T cells were tested against the tetramer^+^ groups, body mass index (BMI), age, and female sex. *p < 0.05; **p < 0.01; ***p < 0.001.

### Systolic blood pressure is associated by trend with higher frequencies of CMV-specific CD45RO^-^/CD27^-^ T_EMRA_

Adding to previous reports on the association of CMV-positivity and risk factors for cardiovascular diseases, we investigated a connection to specific T cells and their differentiation status. 50 participants reported a history of cardiovascular diseases (CVD), including coronary heart disease, myocardial infarction, angina pectoris, heart failure, cardiac arrhythmias and peripheral artery disease. These donors had significantly higher frequencies of CMVpp65-specific CD8^+^ T cells comparing to participants without cardiovascular problems (Figure 5A), and further, systolic blood pressure (SBP) was associated with the overall frequency of these cells (Figure 5A). No association between the CMV-specific T cell frequency and the diastolic blood pressure were detectable (data not shown). Broken down into the subtypes, the correlation of SBT with effector T cells was stronger compared to the overall CMVpp65-specific CD8^+^ T cells, while the naive, T_CM_ and T_TM_ subtypes followed the opposite trend (Figure 5B). Dissecting the effector T further, no significant correlation was observable, although it appears as if mostly the CD45RO^-^CD27^-^ T cells, but not the CD45RO^+^CD27^-^, were increased with higher SBP (Figure 5C). Of note, SBP was also positively correlated with age (Supplemental Figure 5), however, age was associated with the CD45RO^+^CD27^-^ but not the CD45RO^-^CD27^-^ T cells in our hands (Figure 2D).

**Figure 5.**
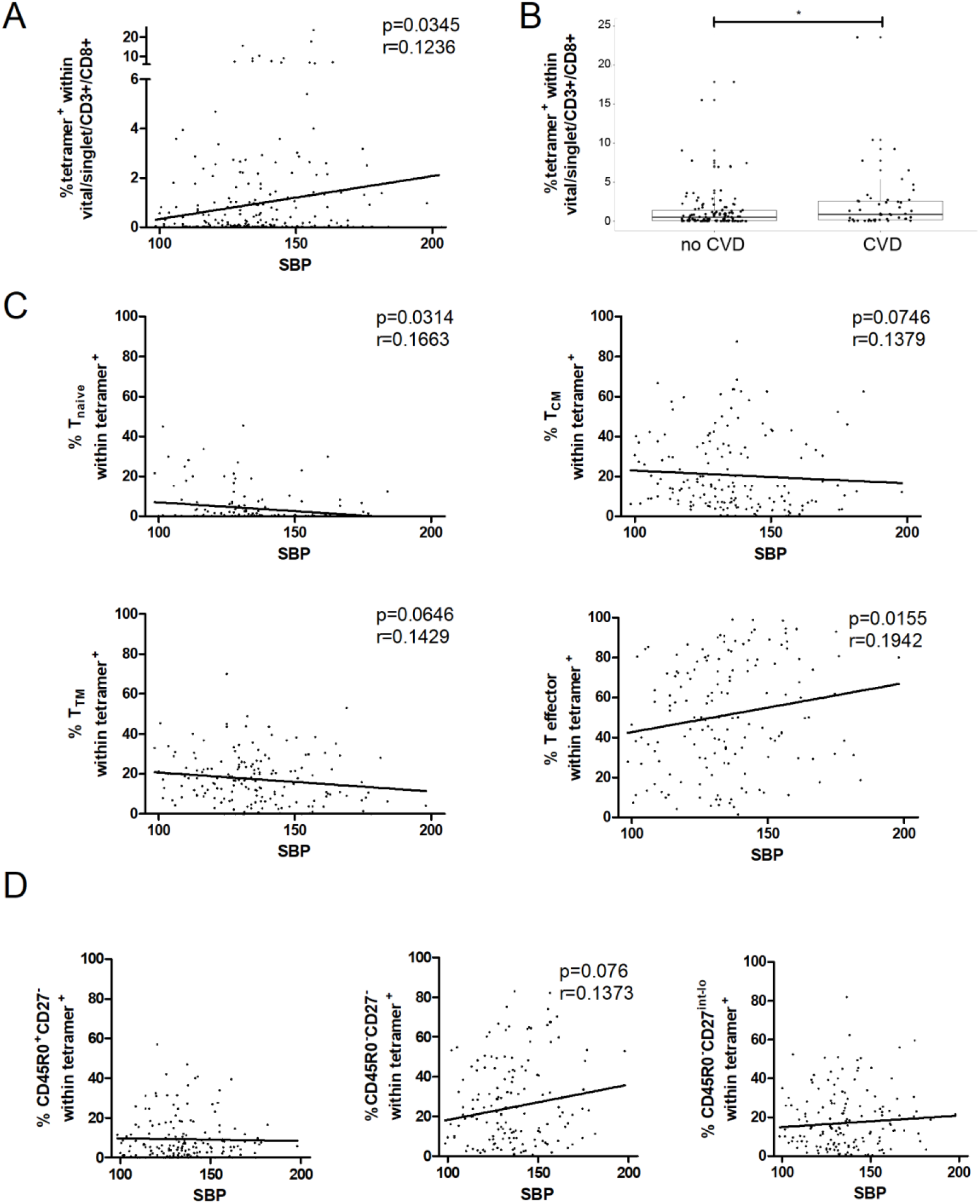
Association of the frequency of CMV-specific T cells and systolic blood pressure. A, Frequency of CMVpp65_495-503_ MHC-I-tetramer^+^ T cells plotted against systolic blood pressure (SBT). B, Frequency of CMVpp65_495-503_ MHC-I-tetramer^+^ T cells comparing individuals with or without a self-reported a history of a history of cardiovascular diseases (CVD), i.e. coronary heart disease, myocardial infarction, angina pectoris, heart failure, cardiac arrhythmias or peripheral artery disease. Wilcoxon matched pairs test, *p < 0.05.C, Frequency of naïve, T_CM_, T_TM_, and effector T cells among CMVpp65_495-503_ MHC-I-tetramer^+^ T cells plotted against SBT. D, CMVpp65_495-503_ MHC-I-tetramer^+^ T effector T cells were further dissected into T_EM_, CD27^-^ T_EMRA_ and CD27^int^ T_EMRA_ groups as indicated. Spearman’s rank correlation, Spearman’s r and p as indicated where applicable.

## Discussion

CMV infection currently affects approximately 56% of people in Germany [22], while in other parts of the world often higher rates are observed [23]. It has been described as one of the most influential non-genetic impacts on the immune system [7], since the reshaping the cellular composition is impacting vaccinations and the immune response to infections [24]. The effects on the immune cell populations are often so strong that they are detectable when analyzing whole blood, not just when focusing on virus-specific T cells. In our cohort, we were able to describe these phenotypically differences by a comprehensive set of high-dimensional spectral flow cytometry distinguishing 97 immune cell types [25]: CMV-seropositivity was associated with increased frequencies of three late-stage CD8^+^/CD27^-^/CD28^-^ T effector cell subtypes, while 16 different T cell clusters, all expressing CD27, were negatively associated. Within the CD4^+^ T cell fraction, four different CD27^-^/CD28^-^ T effector cell clusters were associated with CMV-seropositivity, while all naïve clusters displayed negative associations. The increase in the late-stage effector lymphocyte populations is seen as a result of the continuously attempts to control CMV reactivation and are part of the success of this operation [2, 26].

While many population or cohort studies rely on the CMV-IgG serostatus to investigate the impact of CMV on the immune response, this study used the specific CD8^+^ T cell response as a basis for investigating the remodeling of human immunity by CMV. Thereby we can confirm here most of the earlier findings, including that chronic encounter of the virus leads to accumulation of late-differentiated CD8^+^ T cells. These changes were that strong, that differences were detectable on the level of leukocyte counts in our cohort: total CD8^+^ as well as CD4^+^CD8^+^ double-positive T cell frequencies and absolute counts were increased in individuals with a strong cellular immune response to CMV, independent of age. Earlier studies have shed light on the rare phenotype of CD4^+^CD8^+^ double-positive T cells, which are often found in patients with chronic virus infections [27-30].

Previous studies addressed the number, cytokine production, and growth potential of CMVpp65_495-503_-specific T cells, pointing out a T_EMRA_ phenotype with reduced proliferative capacity [31]. Adding to that, our study shows that different subsets of effector cells are associated with age, seropositivity, and SBT. Especially the CD45RO^+^CD27^-^ T_EM_ and CD45RO^-^CD27^int-lo^ T_EMRA_ fractions of the CD8^+^ T effector cells were increased with age. On the other hand, CMV-seropositivity was merely associated with CD45RO^+^CD27^-^ T_EM_ and CD45RO^-^CD27^-^ T_EMRA_ frequencies, which was mirrored by an increased SBT. This indicates that not age per se renders elderly CMV-positive individuals susceptible to CVD, but the immune response to CMV.

While studies have reported, that a strong response to CMV was associated with longer lifespan [32], the continuous combating the CMV attempts to reactivate is nevertheless associated with certain costs for the organism. The iAge score, which was developed as a predictive score for inflammaging and was shown to be able to predict multimorbidity, correlated with CMV positivity in a multiple regression model [33]. Elderly individuals seropositive for CMV or with clonal expansion of T_EMRA_ cells were reported to show poor humoral responsiveness following influenza vaccination [8, 34-36]. Further, the specific T-cellular response to another herpesvirus, EBV, was reduced in CMV-seropositive compared to seronegative elderly (but not young) individuals [37]. The increased risk of seropositive individuals to develop CVD has been observed decades ago [11], and more recent studies observed that CMV-specific CD8^+^ T cells, as determined by production of cytokines and/or surface expression of degranulation markers after specific stimulation, are correlated with arterial stiffness [38]. Analyzing 207 patients with hypertension, a correlation of arterial stiffness with CMV pp65-specific CD8^+^ T-cell responses could be observed [39]. In our hands it became apparent, that especially the specific T effector-like T cell subset and further the CD45RO^-^CD27^-^ T_EMRA_ cells were increased with SBP. These data suggest that the differentiation status of CMV-specific T cells influences the cardiovascular system, and can serve as a predictor for disease. While aging itself represents a risk factor for hypertension, and overall CD8^+^CD27^-^ T cell frequencies have been reported to predict for this [21], we observed that different subtypes of CMV-specific T cells are associated with aging and SBT, respectively.

It should be noted, that some donors may have been included into the HLA-A*02:01 tetramer-staining that carried other HLA-A2 alleles, since the HLA-type was determined by antibody testing. However, this approach had the advantage of a quick turnover and allowed analyzing the CMV-specific T cells directly without further freezing and storage.

In conclusion, we made use of the RESIST SI cohort, which resembles the aging population in Germany [17], to investigate associations of the specific cellular immune response to CMV with aging and comorbidities. While confirming previous findings, we show that aging is associated with a different phenotype of CMV-specific T cell response compared to the humoral response and an increased SBT. Analysis of specific T cell subtypes may therefore help to predict cardiovascular anormalities.

## Acknowledgements

We express especially thanks to Gabriele Begemann and Petra Kienlin for continuous and expert cell handling and measurements. The generation of the RESIST SI cohort was a collaborative effort of the SI cohort investigator group within the EXC RESIST (https://www.resist-cluster.de/en/), and we thank in this regard especially Jana Heise, Felix Jenniches, Yvonne Kemmling, Berit Lange, Stefanie Castell, Helmholtz Centre for Infection Research (HZI), Braunschweig, Germany; Lennart Riemann, Verena Kopfnagel, Norman Klopp, Thomas Illig, Hannover Medical School (MHH), Hannover, Germany; Nienke van Unen, Xun Jiang, Manoj K. Gupta, Yang Li, Department of Computational Biology of Individualised Medicine, Centre for Individualised Infection Medicine (CiiM), Hannover, Germany. We further wish to thank all the participants and patients who volunteered for this study.

## Author contributions

Study design: L.R., T.W. Experiments: L.R. Data analysis: L.R., S.T. Data interpretation: L.R., B.B., R.F., J.H., T.W. Writing of the first draft: L.R. Editing and revising: all authors. Supervision: R.F., T.W.

## Additional information

Supplementary information is available for this paper. Correspondence and requests for materials should be addressed to Thomas Werfel or Lennart Roesner.

