## Supplement for "Different CMV-specific effector T cell subtypes are associated with age, CMV serostatus, and increased systolic blood pressure"

**Supplemental Data**

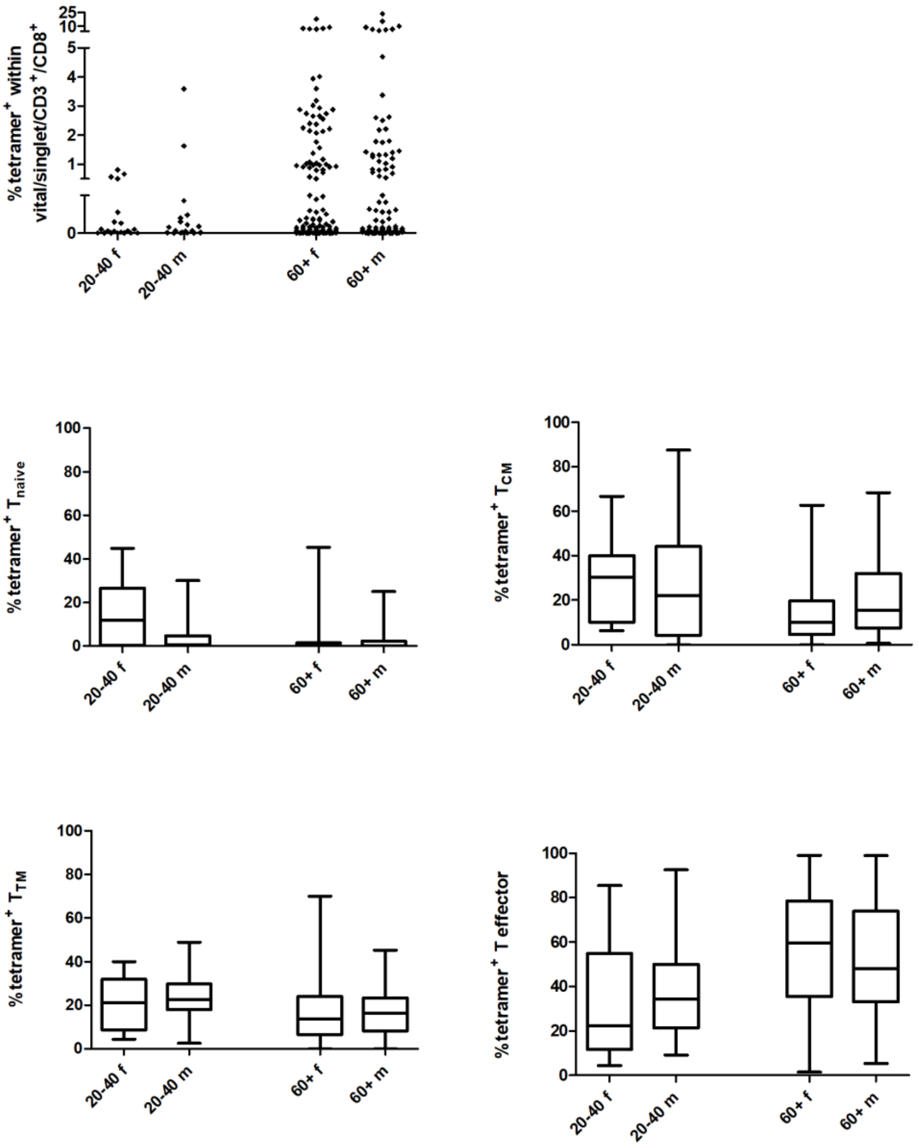


Supplemental Figure 1. **Frequency and differentiation status of CMV-specific T cells with regard to sex.** Upper: Frequency of $CMVpp65_{495-503}$ MHC-I-tetramer$^{+}$ T cells in healthy volunteers by male and female sex. Lower: Frequency of naïve, TCM, TTM, and effector t cells among $CMVpp65_{495-503}$ MHC-I-tetramer$^{+}$ T cells in healthy volunteers by male and female sex.

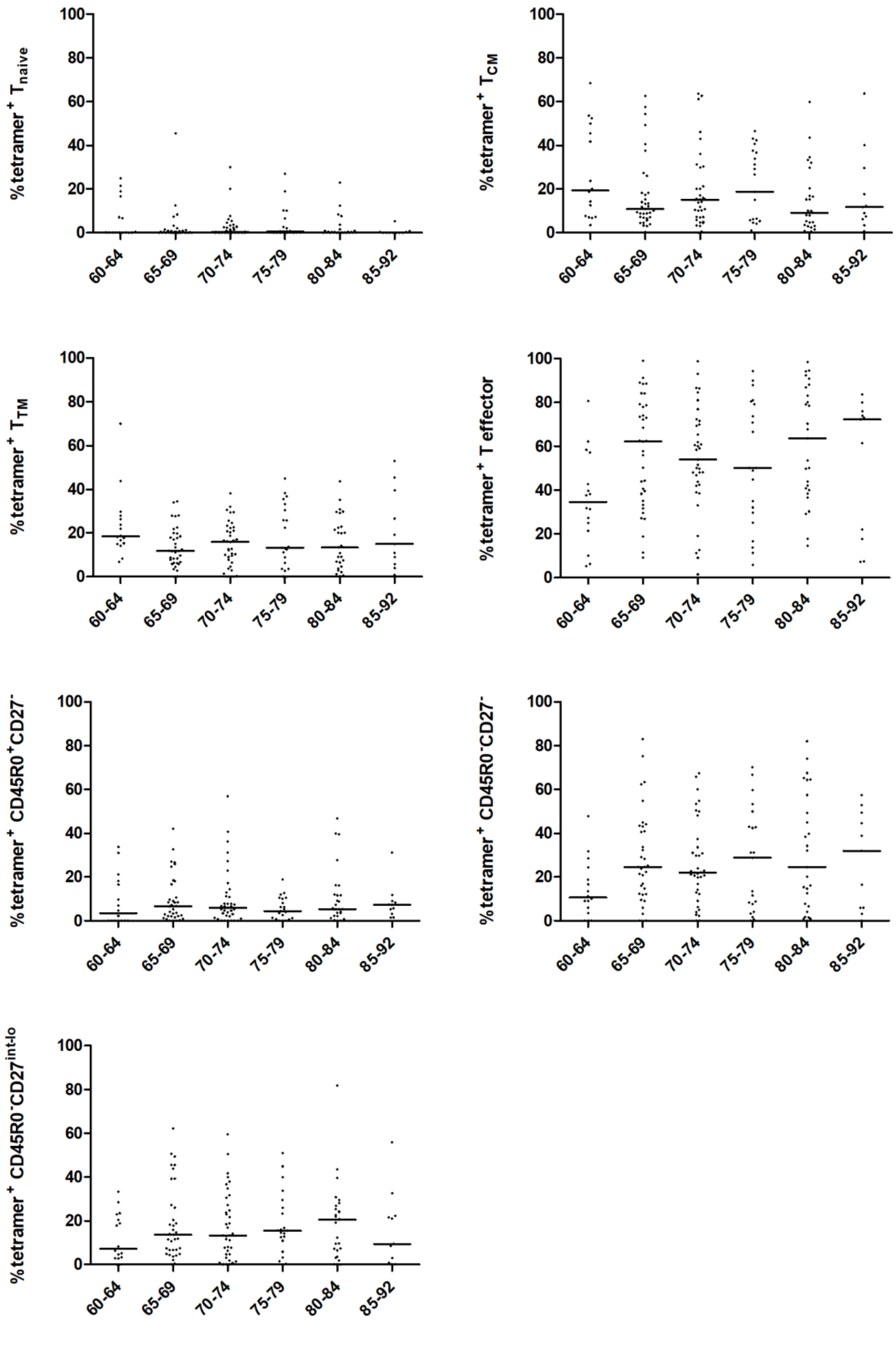


Supplemental Figure 2. **Frequency of naïve, TCM, TTM, and effector T cells among CMVpp65$_{495\text{-}503}$ MHC-I-tetramer$^+$ T cells among the 60+ year old healthy volunteers, dissected into groups of 5 years of age.** CMVpp65$_{495\text{-}503}$ MHC-I-tetramer$^+$ T effector T cells were further dissected into $T_{EM}$, CD27$^-$ $T_{EMRA}$ and CD27$^{int}$ $T_{EMRA}$ groups as indicated.

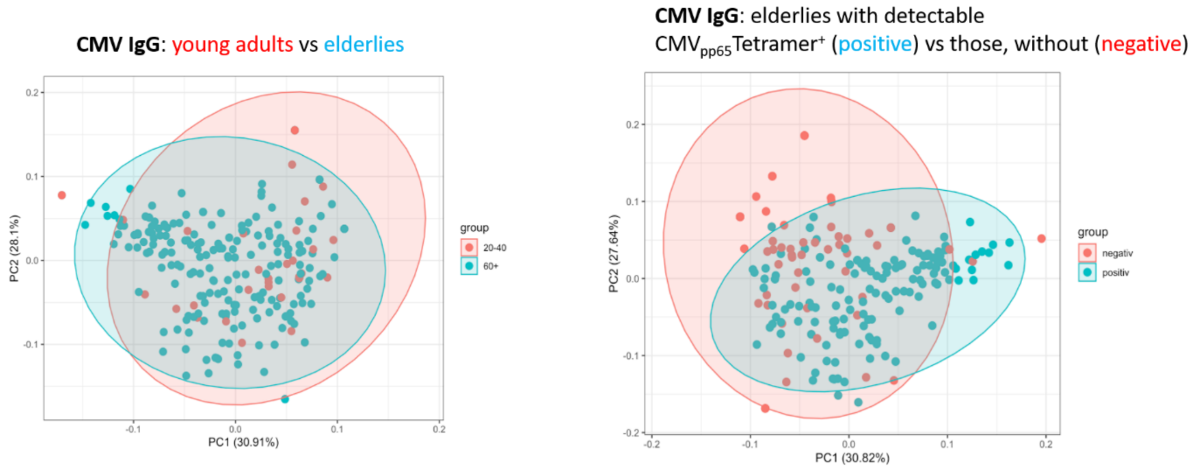


Supplemental Figure 3. **Principal component analyses on anti-CMV IgG serum titers and age groups**. Left: 20-40 year old young adults (red) and 60+ year-olds (blue). Right: 60+ year-olds with (blue) and without (red) detectable CMVpp65$_{495\text{-}503}$ MHC-I-tetramer$^{+}$ T cells.

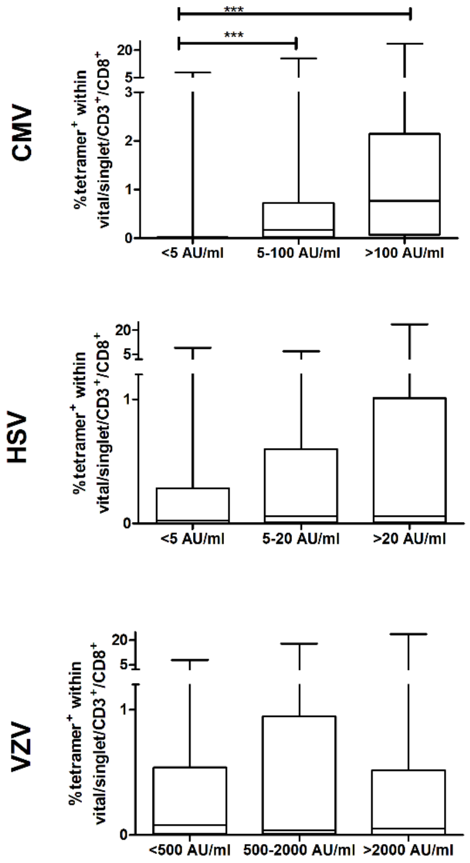


Supplemental Figure 4. **Frequency of CMV-specific T cells with regard to anti-herpesvirus IgG titers.** Frequency of CMVpp65$_{495\text{-}503}$ MHC-I-tetramer$^{+}$ T cells compared among individuals with low, intermediated, and high serum titers against CMV, HSV, and VZV as indicated. Kruskal-Wallis test with Dunn´s multiple comparison test, ***p < 0.001.

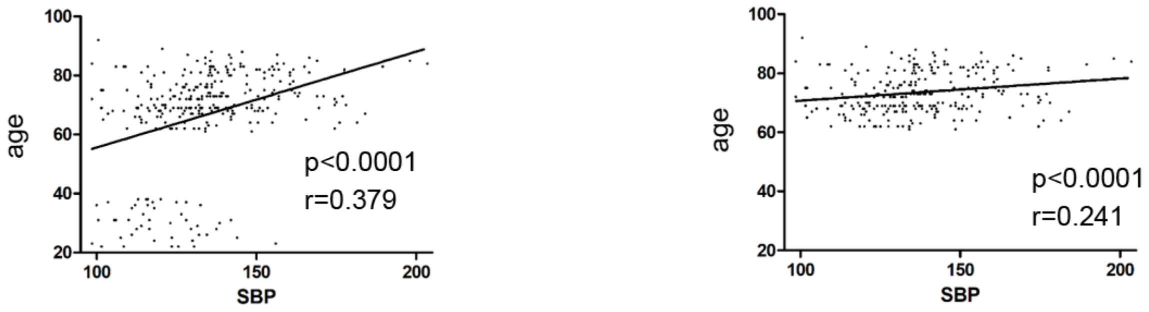


Supplemental Figure 5. **Correlation analysis of age and systolic blood pressure** (SBP). Left, including 20-40 year-olds and 60+ year-olds. Right, Separate analysis of 60-year-olds. Spearman's rank correlation, Spearman´s r and p as indicated.
